# Mechanical mapping of mammalian follicle development using Brillouin microscopy

**DOI:** 10.1101/2021.02.21.432113

**Authors:** Chii Jou Chan, Carlo Bevilacqua, Robert Prevedel

**Affiliations:** Cell Biology and Biophysics Unit, European Molecular Biology Laboratory, Heidelberg, Germany; Mechanobiology Institute, National University of Singapore, Singapore; Department of Biological Sciences, National University of Singapore, Singapore; Collaboration for joint PhD degree between EMBL and Heidelberg University, Faculty of Biosciences.; Developmental Biology Unit, European Molecular Biology Laboratory, Heidelberg, Germany; Epigenetics and Neurobiology Unit, European Molecular Biology Laboratory, Monterotondo, Italy; Molecular Medicine Partnership Unit (MMPU), European Molecular Biology Laboratory, Heidelberg, Germany

## Abstract

In early mammalian development, the maturation of follicles containing the immature oocytes is an important biological process as the functional oocytes provide the bulk genetic and cytoplasmic materials for successful reproduction. Despite recent work demonstrating the regulatory role of mechanical stress in oocyte growth, quantitative studies of ovarian mechanical properties remain lacking both *in vivo* and *ex vivo*. In this work, we quantify the material properties of ooplasm, follicles and connective tissues in intact mouse ovaries at distinct stages of follicle development using Brillouin microscopy, a non-invasive tool to probe mechanics in three-dimensional (3D) tissues. We find that the ovarian cortex and its interior stroma have distinct material properties associated with extracellular matrix deposition, and that intra-follicular mechanical compartments emerge during follicle maturation. Our work provides a novel approach to study the role of mechanics in follicle morphogenesis and pave the way for future understanding of mechanotransduction in reproductive biology, with potential implications for infertility diagnosis and treatment.

---

In mammalian development, folliculogenesis describes the progression of a number of small primordial follicles into large preovulatory follicles with functional oocytes in the ovaries (Fig. 1a) Upon activation, the primordial follicle transitions into primary state, when the surrounding somatic cells (granulosa cells (GCs)) become cuboidal and undergo extensive proliferation. It then develops into a secondary follicle with multiple layers of GCs, basal lamina and a theca layer. This is followed by the formation of fluid-filled antral follicle, and ovulation where the oocyte is released from the ovary. While hormonal signaling is known to impact antral follicle formation onwards, early stages of preantral follicle development are known to rely on intra-follicular signaling^1^. Recently there is emerging evidence that mechanical stress imposed by the extracellular matrix (ECM) plays a role in the oocyte development, such as the activation of primordial follicles^2,3^. Furthermore, it has long been hypothesized that the ovarian cortex and the inner medulla have distinct stiffness^4^, mostly because appropriate tools for quantifying tissue material properties *in vivo* do not yet exist or face important limitations. Currently, Atomic Force Microscopy (AFM)^5^ and micropipette aspiration^6,7^ are most often used to infer mechanical properties, such as elasticity or viscosity in the micrometer regime at cellular surfaces or in superficial tissues. A recent AFM study reported regional difference in ovarian stiffness^8^, although the work was carried out in bisected ovaries which may release tissue stress and perturb the mechanical properties of the ovaries^9^. Other approaches such as optical coherence elastography and stress sensors involving liquid droplets and deformable gel beads have also been developed to infer tissue mechanical properties and stress in 3D^10–14^. However, all these methods are invasive which could perturb the sample, rely on mechanical models to extract local elastic parameters, and are unable to provide a high-resolution spatial map of the measured mechanics.

**Figure 1:**
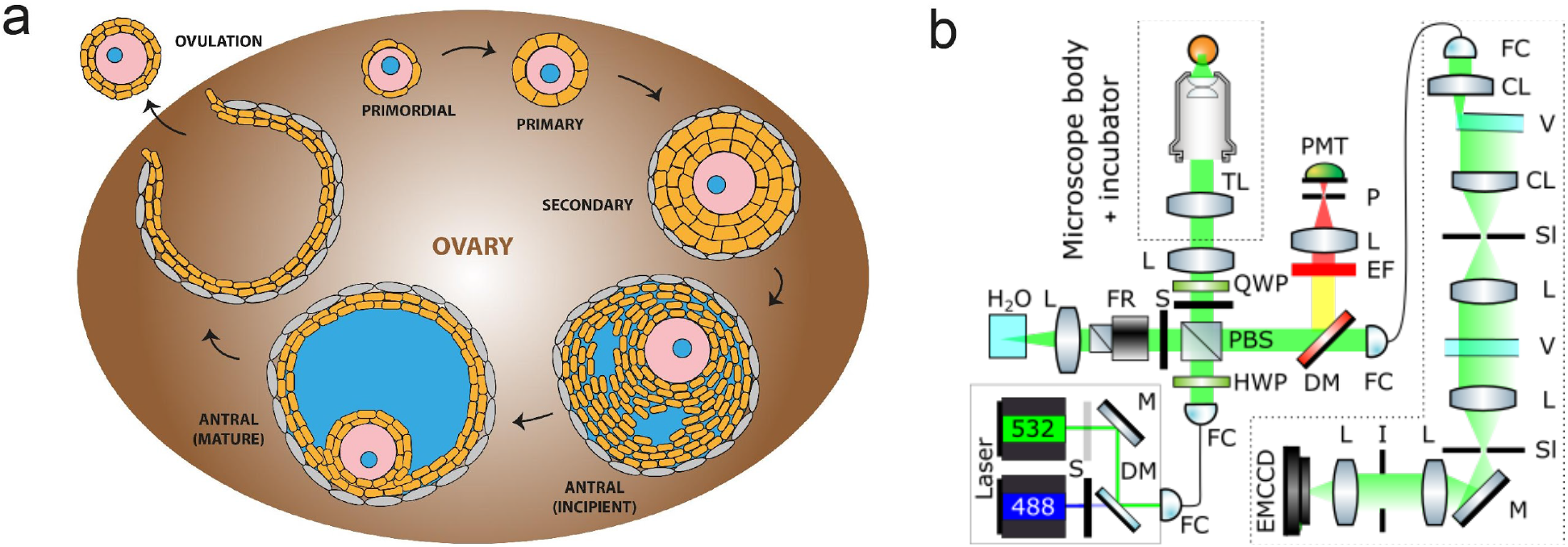
Characterizing mechanical properties of ovaries *ex vivo* with Brillouin microscopy. **a.** Schematic of the developmental cycle of follicles during mouse folliculogenesis. A follicle consists of the oocyte (pink) with its nucleus (blue), surrounded by the somatic cells (orange) and theca cells (grey). The oocyte grows in size during transition to secondary follicle stage, followed by the emergence of fluid-filled lumen (blue) in the antral follicle stage. The oocyte is eventually released during ovulation, and upon fertilization undergoes embryo development. The interstitial tissues of the ovaries comprise of stromal cells, extracellular matrix and vasculature. **b.** Schematic of the confocal Brillouin imaging setup. L=lens, TL=tube lens, PMT=photomultiplier, P=pinhole, EF=emission filter, DM=dichroic mirror, FC=fiber coupler/collimator, QWP=quarter waveplate, HWP=half waveplate, PBS=polarising beam splitter, S=shutter, FR=Faraday rotator, CL=cylindrical lens, V=VIPA, Sl=adjustable slit, V=VIPA, I=iris.

Here, Brillouin microscopy offers an optical and thus non-invasive approach to infer mechanical properties of cells and tissues at high spatial resolution in 3D. It relies on the interaction and inelastic scattering of monochromatic laser light from thermally driven acoustic phonons at high frequencies^15–17^. The scattered light spectrum is indicative of the material’s sound velocity and thus local mechanical properties. In particular, it gives access to the (micro-)elasticity and viscosity through the measurement of the longitudinal modulus and acoustic wave attenuation at the micrometer-scale. Brillouin microscopy is an emerging technique that has been successfully applied to questions in cell biology^18,19^, development^20–22^ as well as within a medical context^23,24^. These studies have typically compared tissue stiffness based on the shift of the Brillouin spectrum alone. While this has become common and widespread practice in the field, the Brillouin shift is also dependent on the material’s refractive index and mass density. Therefore, in situations where a homogenous refractive index cannot be assumed, it is thus impossible to separate the contributions of mechanical properties and the refractive index or density to any observed spatial or temporal changes in the Brillouin frequency shifts. To address this intrinsic limitation, in this study we focus on a less ambiguous metric, the Brillouin loss tangent (BLT), which allows us to single out the mechanical contribution to the observed Brillouin signals.

In this study, we examined the mechanical properties of the mouse ovarian cortex and interior using confocal Brillouin microscopy, focusing on the extra- and intra-follicular tissues during follicle development. Our results demonstrate clear changes in the mechanical environment that the follicles experience during development, which correlate with follicular cell differentiation and extracellular matrix deposition during folliculogenesis.

## Results

### Quantifying cell and tissue material properties with Brillouin loss tangent (BLT)

To address the above-mentioned technical limitation with respect to characterising mechanical properties in 3D, we utilized Brillouin microscopy, which is based on high-resolution spectroscopic measurements of the light scattered by the material under investigation (see **Fig. 1b** and **Methods**). In particular, the backscattered light spectrum, i.e. its peak position and linewidth, provides information about the longitudinal modulus at hypersonic (GHz) frequencies. From the spectrum’s frequency shift, ν, and its full-width at half-maximum (FWHM), Γ, the storage and loss moduli of the sample can be inferred as M’ = ϱ(νλ/2*n*)^2^, and M’’ = ϱΓν(λ/2*n*)^2^, which accounts for the elastic and viscous behaviour of the sample^16^. Here, ϱ is the density and *n* the material refractive index within the scattering volume, and λ the incident wavelength. As is evident from their definition, both the local refractive index and density are central to a proper quantification of the mechanical properties. However, techniques such as tomographic phase microscopy^25,26^ or optical diffraction tomography^27–29^ that can provide information about optical properties in three dimensions and *in situ*, do not work in thick and highly scattering tissues such as the ovary.

In order to decouple the effect of mechanical and optical properties in Brillouin microscopy, here we utilize the so-called Brillouin loss tangent (BLT), defined as tan(ϕ) = M’’/M’ = Γ/ν. By its definition, the BLT does not depend on the sample refractive index and density and thus provides a simple approach to determine whether mechanical properties are the main contributor to observed changes in the Brillouin spectrum and their spatial maps. A similar approach has been recently used to study bulk mechanical properties and their differences in tumor spheroids^24^ and cellulose fibers^30^, but has not yet been used for spatial mapping. From a mechanical viewpoint, the loss tangent can be understood as the relative contribution of effective viscosity to longitudinal modulus within the probed, micrometer-scale region of the sample. It is therefore a measure of how well energy is stored or dissipated in the tissue. In general, a high loss tangent indicates a more liquid-like behaviour and therefore a more pronounced energy dissipation of the (local) phonons^31^. On the contrary, a small loss tangent reflects a dominant solid-like elastic behaviour.

To validate the loss tangent as a meaningful parameter in comparing mechanical properties of biological samples, we imaged and compared the Brillouin frequency shift, linewidth and BLT of the nucleoli and nucleoplasm in mouse embryos at the preimplantation stage. As shown in **Supplementary Fig. 1d**, the nucleoli have significantly higher BLT compared to the nucleoplasm, suggesting that the nucleoli contain more dissipative elements than the nucleoplasm at the microscopic scale. This is consistent with the emerging evidence that the nucleoli are membraneless organelles that exhibit liquid-like properties such as fusion and phase separation in diverse biological contexts^32–34^ According to the above definition of the BLT, this also implies that the phonons experience higher dissipation in the nucleoli.

### The ovary exhibits regional difference in tissue mechanical properties

We first investigated if there exists any regional difference in the ovarian mechanical properties during folliculogenesis (**Fig. 2a,b**). While Brillouin shift data showed variations in the signal intensities across the tissue that likely correspond to the different structural components, shift alone is not sufficient to inform mechanical variations due to its dependence on optical properties, as discussed above. We therefore extracted the BLT from the spectral data, which showed a visibly higher signal in the P7 and P14 ovarian interior compared to the cortices where the primordial follicles typically reside. This suggests that the ovarian cortex may be more elastic than the ovarian interior, which is populated with the larger secondary follicles and stroma that consist of collagen fibres, fibroblasts and vasculature^35^. Of note, this regional difference in BLT arises mainly from a change in the Brillouin linewidth (**Supplementary Fig. 2**), suggesting that the ovarian mechanics is largely dominated by changes in tissue micro-viscosity. Zooming into the primordial follicles, we observed that they are characterised by oocytes with a low BLT, surrounded by a somatic cell layer with higher BLT (**Fig. 2c).** Notably, we also observed a ring of significantly higher BLT, possibly associated with the presence of basement membrane that encapsulates the primordial follicles^1^ (**Fig. 2d**).

**Figure 2:**
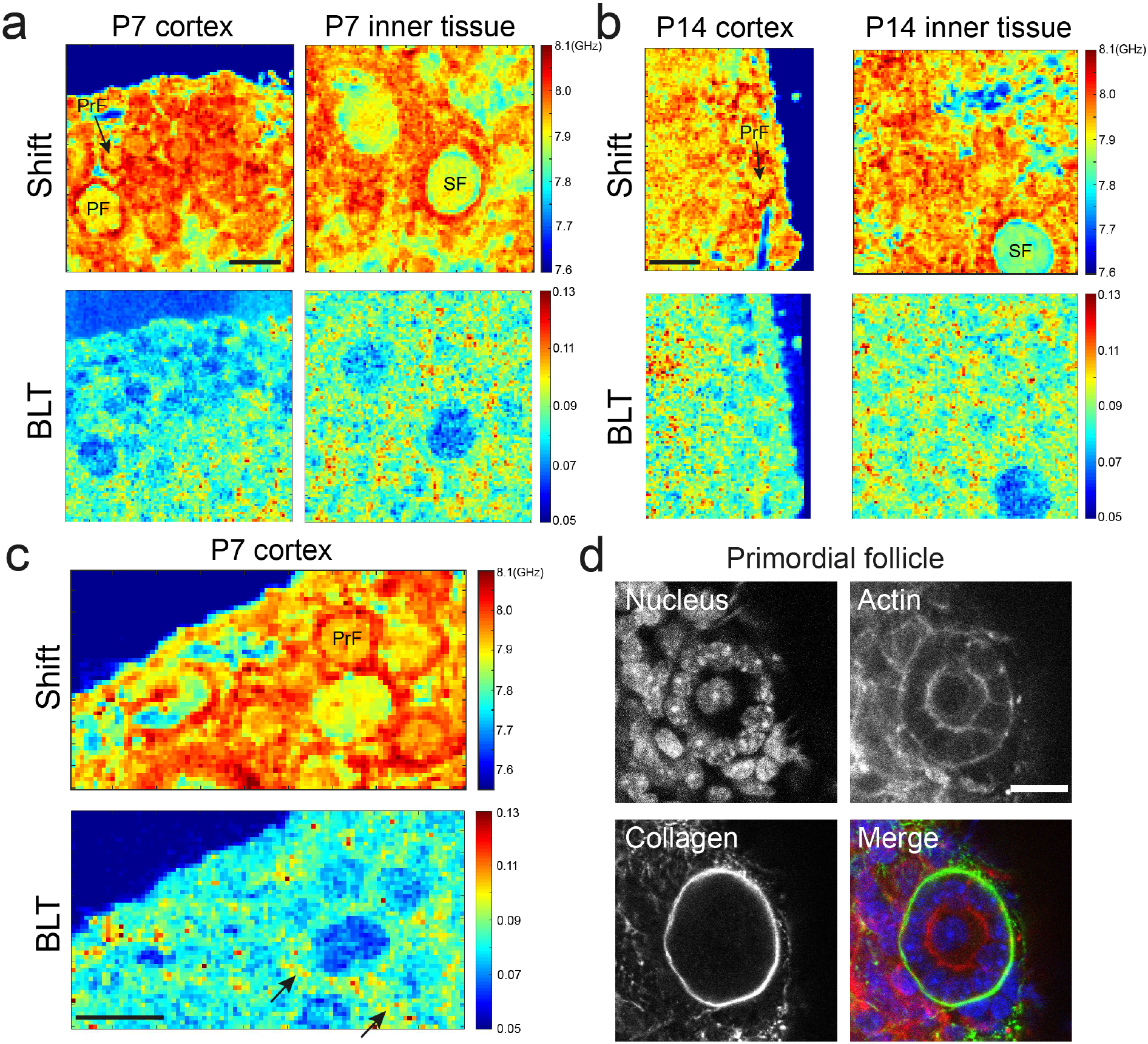
Ovary tissue is characterized by regional differences in mechanical properties. **a.** Maps of Brillouin shift and the corresponding BLT for a P7 ovarian cortex and the inner tissue. Both images were obtained from the same ovary. Primordial follicles (PrF) and primary follicles (PF) reside primarily in the cortex while the secondary follicles (SF) tend to be found in the inner part of the ovary, which shows a higher interstitial BLT. **b.** Maps of Brillouin shift and BLT for a P14 ovarian cortex and the inner tissue. Similar to P7 ovaries, higher PLT is associated with the inner tissue of the ovaries, compared to the cortex. **c.** Zoom in view of P7 ovarian cortex showing higher BLT of cells (black arrows) surrounding the oocyte, which may be associated with the presence of collagen surrounding the primordial follicle, as shown in (**d**). Scale bar = 40 μm in (**a**), (**b**) and (**c**), and 15 μm in (**d**).

### Mechanical compartments emerge in the follicles during follicle maturation

We next performed mechanical mapping of follicles from the secondary to the antral follicles stage, which is marked by the presence of large fluid-filled lumena. Here, remarkably, we found that during this transition an outer tissue ‘shell’ with a significantly higher BLT starts to form and encapsulate the follicle (**Fig. 3b**). This mechanical compartmentalisation becomes most pronounced at the mature follicle stage. The typical width of this ‘shell’ is larger than 10 μm, suggesting that they originate from cellular entities, possibly from the theca cells. This is also consistent with the fact that the theca cells become more established at the antral follicle stage, compared to the secondary follicle stage^36^.

**Figure 3:**
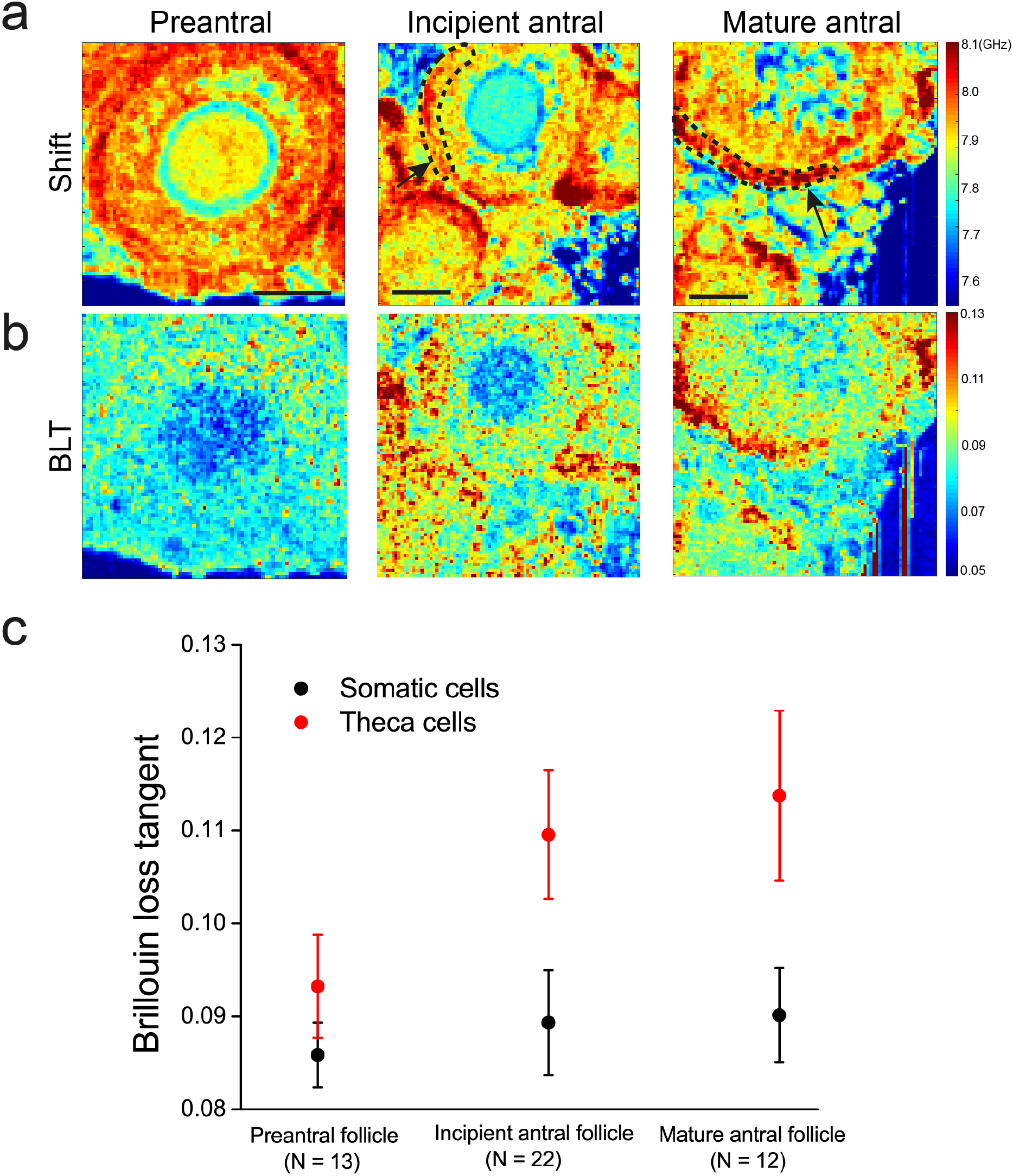
Distinct mechanical compartments emerge during follicle maturation. **a.** Row shows representative maps of Brillouin shift for follicles at distinct stages of folliculogenesis. Scale bar = 40 μm. **b.** Corresponding maps of BLT for **a**. **c.** Plot of BLT for the outer theca cells versus the inner somatic cell layer of follicles at various stages of development. White arrows indicate the follicles under consideration, black dashed lines indicate the region of interest for the theca cell layer.

Of note, Brillouin imaging allows us to visualise the microlumina in the early antral follicles and the larger singly resolved lumena in the late antral follicles (see blue regions within the follicles in the shift maps **Fig. 3a** and **4a**), which is not possible using immunostaining or live dye assay. This ability to image luminal structures in living tissues in a label-free manner based on the different tissue/fluid material properties is a unique strength of Brillouin microscopy. Future longitudinal studies of cultured antral follicles with Brillouin microscopy may reveal key mechanisms of lumen initiation and coalescence during antral follicle morphogenesis.

**Figure 4:**
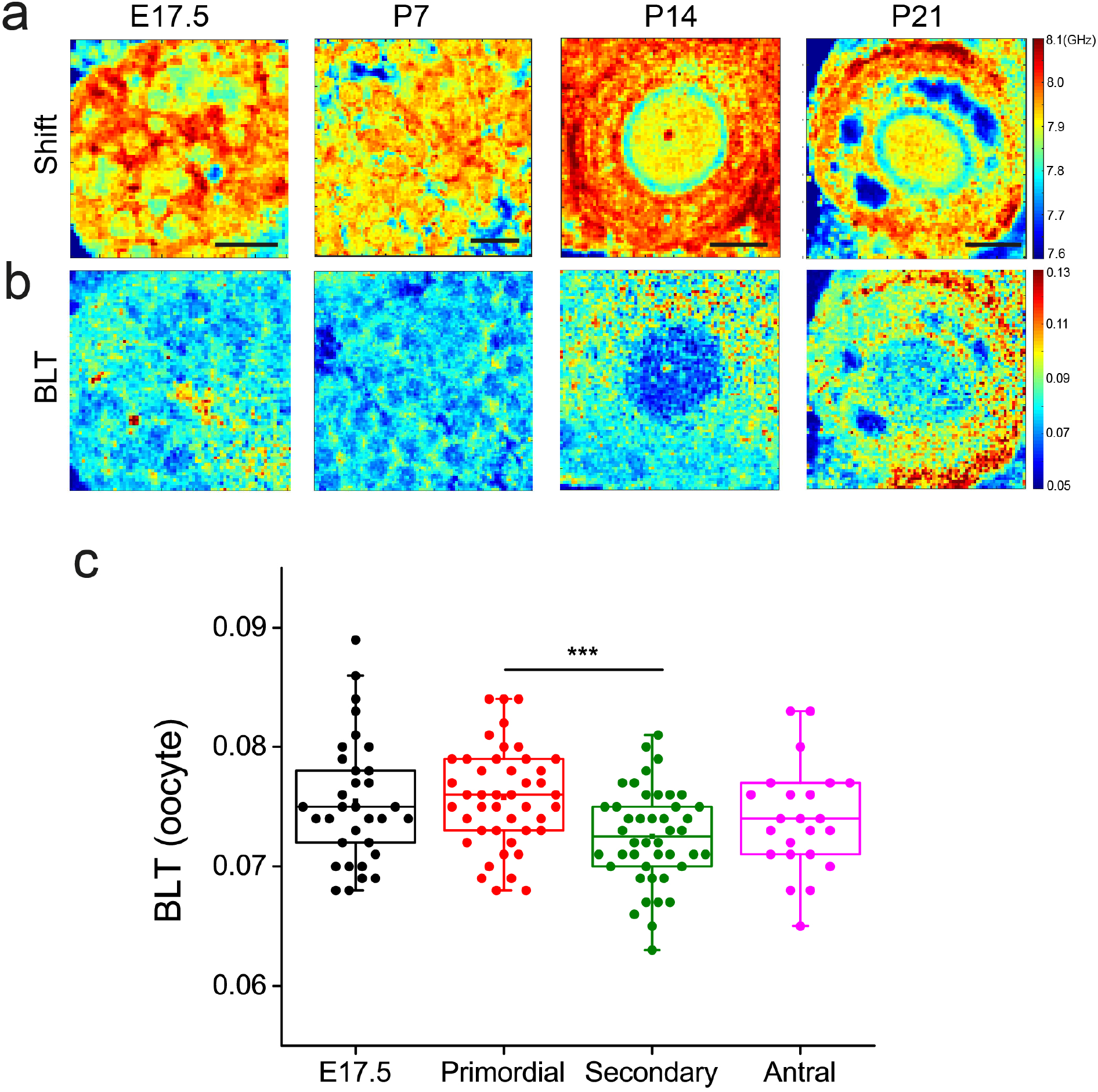
Ooplasm exhibits more liquid-like behavior at the early stages of oogenesis. **a.** Row shows representative maps of Brillouin shift for oocytes from fetal ovaries (E17.5), primordial follicles (P7), secondary follicle (P14) and antral follicle (P21) ovaries. Scale bar = 40 μm. Blue regions within the antral follicle (P21) indicate fluid lumena. **b.** Corresponding Brillouin loss tangent for **a**. **c.** Boxplot of BLT for oocytes at various stages of follicle development. Each data point corresponds to the average signal of one oocyte. N = 35, 45, 45, 21 for E17.5 germ cells and oocytes from primordial, secondary and antral follicle stage, respectively. \*\*\**P*< 0.001.

Following early studies showing changes in the ultrastructure of the oocytes at various stages of growth^37,38^, we next quantify the change in ooplasmic micro-viscoelasticity during folliculogenesis (**Fig. 4**). Interestingly, we observed that the primordial follicles in all stages of follicle development exhibit a small but significantly higher BLT compared to the larger oocytes from the secondary and antral follicles (**Fig. 4c**). Germ cells from the foetal ovaries (E17.5), which are not surrounded by the somatic cells, also have a relatively high BLT compared to the late follicles such as the secondary or antral follicles. These results suggest that the mid- and late stage oocytes are more solid-like at the micro-structural level, compared to the early oocytes.

### Ovarian viscoelasticity is governed by extracellular matrix

To further investigate the origin of changes in BLT signals and ovarian mechanics, we studied the distribution of fibrillar collagen, a known contributor to tissue viscoelasticity^39^, in mouse ovaries at various stages of follicle development. Using Second Harmonic Generation (SHG) imaging, we observed clear interstitial collagen signals in P14 and P21 ovaries but not in P7 ovaries (**Supplementary Fig. 3**). Primordial, secondary and antral follicles are embedded in the collagen network, which became denser and more bundle-liked in the P21 ovaries. This progressive increase in interstitial collagen network during follicle maturation correlates well with the increase in interstitial BLT (**Fig. 2a,b**), suggesting that ECM deposition regulates the ovarian tissue mechanics. To test this hypothesis, we treated the ovaries with CTK, a solution containing collagenase that digest the ECM (see methods). This treatment led to disruption of tissue integrity, as shown by the increased interstitial spacing and disconnected follicles (**Fig. 5a**), as well as an overall qualitative decrease in BLT.

**Figure 5:**
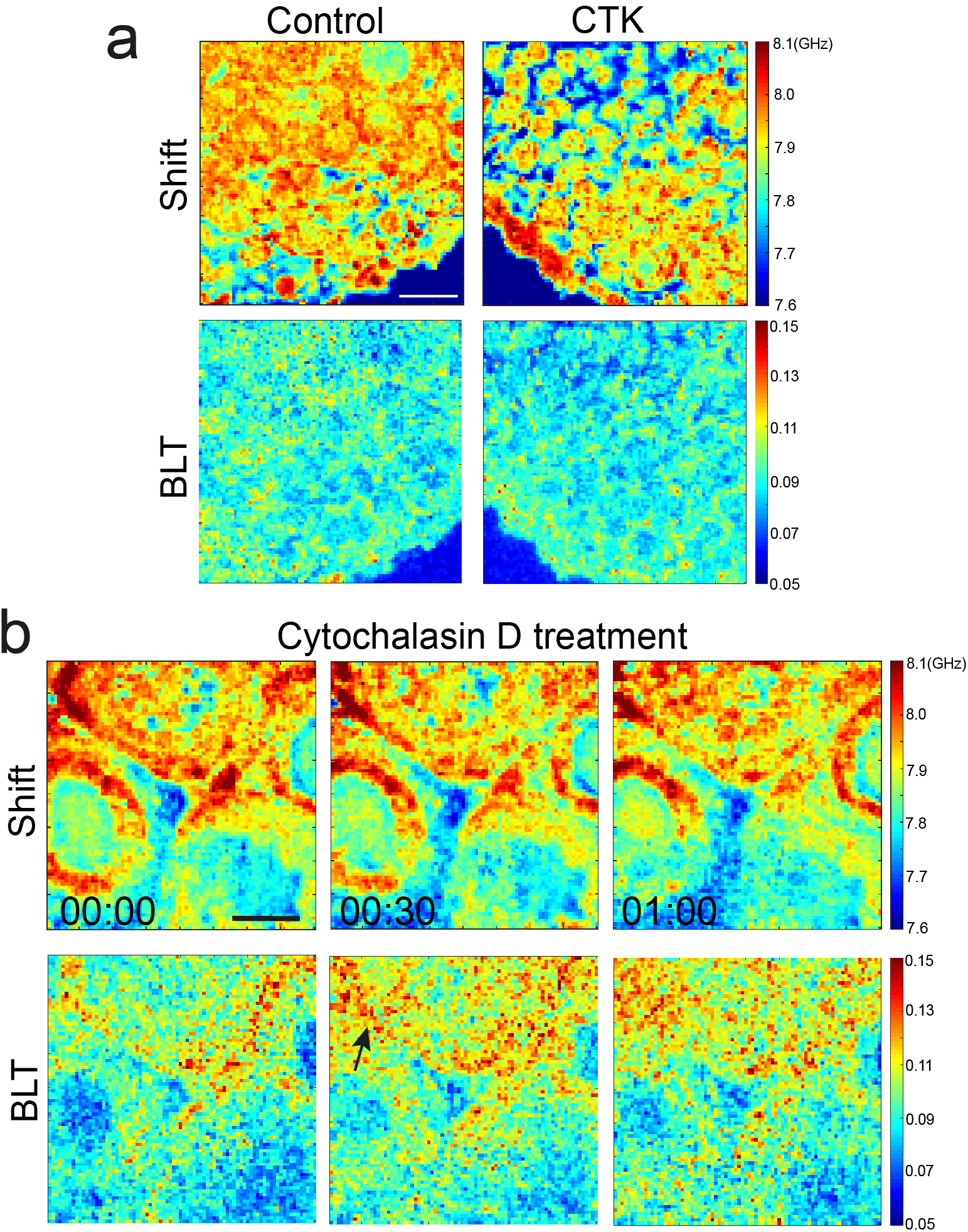
Disruption of extracellular matrix and actin cytoskeleton impacts interstitial viscoelasticity. **a.** Top row: Brillouin shift of a P14 ovary when treated with CTK, which digests collagen matrix. CTK leads to a clear increase in the interstitial space (blue) between the primordial follicles at the cortex. Bottom row: Corresponding BLT maps. Scale bar = 40 μm. **b.** Top row: Brillouin shift of a P14 ovary when treated with cytochalasin D, a pharmacological inhibitor that disrupts actin cytoskeleton. Lower row: Corresponding BLT maps showing that a disruption of actin cytoskeleton leads to higher BLT (black arrow) and decreased elasticity of the interstitial tissues. Time denotes hh:mm.

We further performed immunofluorescence staining, which showed that the actin cytoskeleton exists abundantly within the follicles and the surrounding stroma (**Fig. 2d**). To ascertain the role of actin cytoskeleton in ovarian tissue viscoelasticity, we treated the ovaries with cytochalasin D, an inhibitor of actin polymerization. This led to a partial increase in intra and inter-follicle tissue BLT (**Fig. 5b**), indicating that the ovarian tissue is becoming more liquid-like, consistent with the role of actin cytoskeleton in maintaining cellular elasticity^17^.

## Discussion

In this study, we demonstrated, to our knowledge, the first non-invasive mechanical mapping, with micron-level spatial resolution, of ovarian mechanics *in situ*. From a biological perspective, our work represents a first step towards addressing the mechanical aspects of folliculogenesis, which has gained recognition due to emerging evidence that mechanical stress in the ovary can influence the quality of the growing oocytes through ECM stiffness^40,41^, Hippo signalling pathway^42,43^ and increased fibrosis^44,45^.

It is important to note that Brillouin scattering probes the material properties in the GHz frequency range, which is in contrast to many existing methods currently utilized in biomechanics. This means that the elasticity (longitudinal modulus) measured by Brillouin scattering will typically assume much higher values (in the GPa range) for common biomaterials, compared with the widely used Young’s modulus *E* (often in the kPa range). Nevertheless, we have shown that in this regime, the connective tissues of the ovarian cortex are more elastic compared to the interior of the ovary, which appears more viscous-like (**Fig. 2**) and is enriched with fibrillar collagen (**Supplementary Fig. 3**). Indeed, while there are substantial research to support the elastic role of ECM, the porous nature of the ECM in causing substantial viscous dissipation has only been getting recognized more recently^39^.

Furthermore, the higher BLT associated with the ECM is largely attributed to the variation in the linewidth (**Supplementary Fig. 2)** and the microscopic viscosity, which could arise from hydrophobic hydration around the collagen^17,46^. From the biological perspective, given that the dormant primordial follicles are known to reside primarily in the cortex while the larger secondary follicles are found in the ovarian interior, this raises the intriguing possibility that the regional difference in ECM viscoelasticity may influence the follicle positioning and ovarian tissue patterning

We have shown that the BLT can serve as a useful experimental metric to characterise the 3D tissue material properties in living mouse ovaries. In contrast to the conventionally used Brillouin shift, which is convoluted with the influence of refractive index and mass density, the BLT is independent of these parameters and therefore serves as an unambiguous metric to compare regional differences in tissue viscoelasticity across space and time. While a recent study has combined Brillouin microscopy with optical diffraction tomography to uniquely determine the refractive index and thus quantitative longitudinal modulus of cells on 2D substrates^27,29^, it remains challenging to extend this approach to thick (>100 *μ*m) tissues. In such scenarios, utilizing and quantifying the BLT may be the best alternative to investigate the mechanical properties in a non-invasive and label-free fashion.

At the intra-follicular level, we showed that the periphery of the primordial follicles has a higher BLT associated with the basement membrane (**Fig. 2c**). In view of recent evidence that the basement membrane plays role in maintaining primordial follicle dormancy^1^, our finding suggests that this may involve some mechanotransduction pathways triggered by the distinct basement membrane viscoelasticity. In secondary and antral follicles, the theca cells were found to be more ‘liquid-like’ compared to the somatic cells (**Fig. 3**). This may be due to the fact that the theca cells are associated with increased expression of hyaluronic acid, which is known to promote tissue hydration^45^. The theca cell layer therefore provides a softer microenvironment that may serve as a mechanical ‘sponge’ to buffer the oocyte and somatic cells against external mechanical stress through stress dissipation. The exact origin and functions for the theca cell mechanics will be exciting topics for future studies. The reduction in ooplasmic BLT during the follicles’ transition to the secondary follicle stage suggests that the ooplasm of the oocytes are becoming more solid-like. Interestingly, oocytes are known to undergo dramatic volumetric expansion (~10-fold) during secondary follicle development. Whether this rapid growth in oocyte size leads to ooplasmic dilution and more elastic behavior remains to be investigated in the future, possibly with the use of other techniques such as microrheology^47^.

Overall, our work demonstrates the power of Brillouin microscopy in mechanical phenotyping the different cell types (oocyte, somatic cells and theca cells) in developing follicles. The unique approach of Brillouin microscopy to track cellular dynamics in a label-free and 3D manner harbours great potential for future investigation of the roles of mechanics in reproductive ageing^24^ and ovarian diseases (e.g. polycystic ovary syndrome)^48^.

## Data Availability

Data that support the findings of this study is available from the corresponding authors upon reasonable request.

## Acknowledgements

We are grateful to Alba Diz-Muñoz and the European Molecular Biology Laboratory (EMBL), especially the animal facility, electronic and mechanical workshop for their support. We thank Katsuhiko Hayashi for scientific discussion and training on mouse ovary work, and Takashi Hiiragi for his support in preliminary work. We thank Kareem Elsayad, Saw Thuan Beng and Tetsuya Hiraiwa for critical reading and constructive feedback on the manuscript. The work was supported by the European Molecular Biology Laboratory and an ERC Consolidator Grant to R.P. (no. 864027, Brillouin4Life).

## Author contributions

C.J.C conceived the project, designed the experiments, and wrote the manuscript with input from all authors. C.B. and R.P. designed and set up the Brillouin microscope. C.J.C performed experiments and analysed data, with help from C.B. R.P. supervised, obtained funding and contributed to the writing of the manuscript.

## Methods

### Ovary work

Embryonic and postnatal mice ovaries were obtained in the animal facility at the European Molecular Biology Laboratory, with permission from the institutional veterinarian overseeing the operation (ARC number 2020-01-06RP). The animal facilities are operated according to international animal welfare rules (Federation for Laboratory Animal Science Associations guidelines and recommendations).

### Imaging setup

The setup of the confocal Brillouin microscope has been described previously^25^, and its schematic is depicted in Fig. 1b. To image the ovaries in physiological conditions, the microscope body (Zeiss Axiovert 200M) is equipped with a custom-built incubation chamber that can actively control the temperature, CO_2_ and O_2_ levels and monitor humidity. The sample is positioned on a 3-axes Piezo translational stage (P-545.3R8H, Physik Instrumente) that allows point-scanning with sub-micrometer precision. The laser source for Brillouin imaging is a 532-nm CW single mode laser (Torus, Laser Quantum); The setup includes also a 488-nm diode laser (Omicron Luxx 488-60) used as an excitation source of GFP-like fluorophores. The two lasers are combined, by means of a dichroic mirror, and coupled into the same PM single-mode fiber to ensure collinearity and a clean beam profile at the output. After collimation the light passes through a half waveplate (HWP) followed by a polarising beam splitter (PBS): the rotation of the HWP determines the ratio of intensities of the beams transmitted and reflected by the PBS, that are delivered respectively to the sample and to a cuvette filled with water for calibration. Two shutters, controlled via a self-written LabVIEW software, allow to collect a calibration spectrum every 50 points. The periodic calibration helps correct for the drift of the laser wavelength over time. The laser is focused on the sample by a 1.0 NA oil objective that gives a spatial resolution of 0.30 μm × 0.30 μm × 1.76 μm (FWHM), as measured from the edge response of the Brillouin amplitude at an oil water interface^49^. The backscattered light is collected by the same objective and, after being reflected by the PBS, is coupled into a single mode fiber and delivered to the Brillouin spectrometer. A narrowband bandpass filter (Semrock LL01-532) before the fiber coupler reflects the emitted fluorescence while transmitting the Brillouin signal. The fluorescence signal is detected by a photomultiplier tube (Thorlabs PMT1001) preceded by a pinhole that ensures confocality.

The Brillouin spectrometer is based on a two-stage virtually imaged phased array configuration^23^, featuring two VIPAs with a nominal free spectral range of 30 GHz and spectral resolution of 520 MHz. The addition of a Lyot stop^50^ before the camera (iXon DU897, EMCCD camera; Andor Technology) brings the suppression of elastically scattered light to ~65 dB. The spectral resolution of the spectrometer is comparable to the typical linewidth in biological matter, therefore deconvolution is needed to obtain a sensible value for the linewidth. Since the lineshapes for both the spectral resolution and the Brillouin linewidth are Lorentzian, we performed deconvolution by subtracting the measured spectral resolution (FWHM of the laser) from the linewidth fitted to the experimental spectra. The linewidth is also affected by the use of high NA optics^51^, but we decided not to correct for this effect because of several reasons: the correction would be exact only for isotropic samples; the broadening is affecting all the experimental points in the same way, and the broadening is reduced in the 180 degrees scattering geometry.

### Data analysis and statistics

The acquired scattering spectra were analyzed in real time with a custom-written Labview program by fitting Lorentzian functions to the spectral data. This yielded the position, width and intensity of the Brillouin peaks. Pixel size was chosen to be 2 μm, an optimal compromise balancing resolution and Field-of-view (FOV) which was chosen to be 180 × 180 *μ*m unless otherwise stated. Overall image acquisition time is typically around 15 min. Fitting may be noisy due to low SNR in deep tissues, hence only images or ROIs were selected for subsequent analysis whose datapoints displayed a fit with an *R^2^* greater than 0.92. The images of Brillouin shift and linewidth were smoothened with Gaussian filter (radius 0.5 pixels, ImageJ), and the BLT data were generated using MATLAB. ROI for extra-follicular tissues, oocytes, somatic cells and theca cells were selected by checking co-localisation of the BLT and Brillouin shift data, which showed a better contrast for these cellular structures.

All statistical analysis was performed using the software Origin (version 8.5.6; OriginLab, Northampton, MA). Significances for data displaying normal distributions were calculated with unpaired Student’s *t*-test (two-tailed, unequal variance). All box plots extend from the 25^th^ to 75^th^ percentiles (horizontal box), with a line at the median and whiskers extending to max/min data points.

### Second Harmonic Generation imaging and analysis

Ovaries from P7, P14 mice, and mice older than P21 were harvested and placed in a glass slide with conical well, filled with 200 μl of PBS and covered with #1.5 coverslip (Thermo Fisher). SHG imaging of mouse ovaries was performed on Zeiss NLO LSM780 microscope equipped with a fs-pulsed laser. Ovaries were excited with a wavelength of 800 nm through a 25X (0.8 NA, water) objective. SHG signals from the tissues were captured in the backward direction in the autofluorescence channels of the microscope.

### Immunofluorescence staining

Ovaries were fixed with 4% PFA at 4°C for 1 hr. Following washing with PBS (1 hr), samples were placed in blocking solution (PBS + 0.1% BSA (Sigma, 9647) + 0.3% Triton X-100 (Sigma, T8787)) and kept. Samples are then placed in blocking solution added with primary antibody and kept at 4°C overnight. The samples were then washed three times with PBST (PBS + 0.1% Tween 20) for 1 hr each time, before being kept at 4°C overnight in secondary antibody diluted in blocking solution. Phalloidin and DAPI diluted 1:1000 in Blocking One Histo were added at this stage. Ovaries were then washed three times in PBST for 1 hr each time, before being mounted in Fluoro-KEEPER antifade reagent (Nacalai) followed by imaging.

Primary antibody against collagen IV (Millipore, AB756P) was used at 1:100. Secondary antibodies targeting rabbit immunoglobulin-coupled Alexa Fluor 546 (Invitrogen, A10040) was used at 1:200. Alexa Fluor 633-coupled phalloidin (Invitrogen, R415) was used at 1:50. DAPI (Invitrogen, D3751) was used at 1:1000.

### Pharmacological inhibition

Following a similar protocol as a recent study^2^, CTK reagent was prepared as 1 mM CaCl_2_, collagenase type IV (0.1 mg/ml), 20% KSR (Invitrogen), and 0.025% trypsin EDTA (Invitrogen). Ovaries were treated with PBS or CTK for 1 hour at 37°C in a CO_2_ incubator before imaging. Cytochalasin D (Sigma) was used at a final concentration of 20 μM.

## Supplementary Information

**Supplementary Figure 1:**
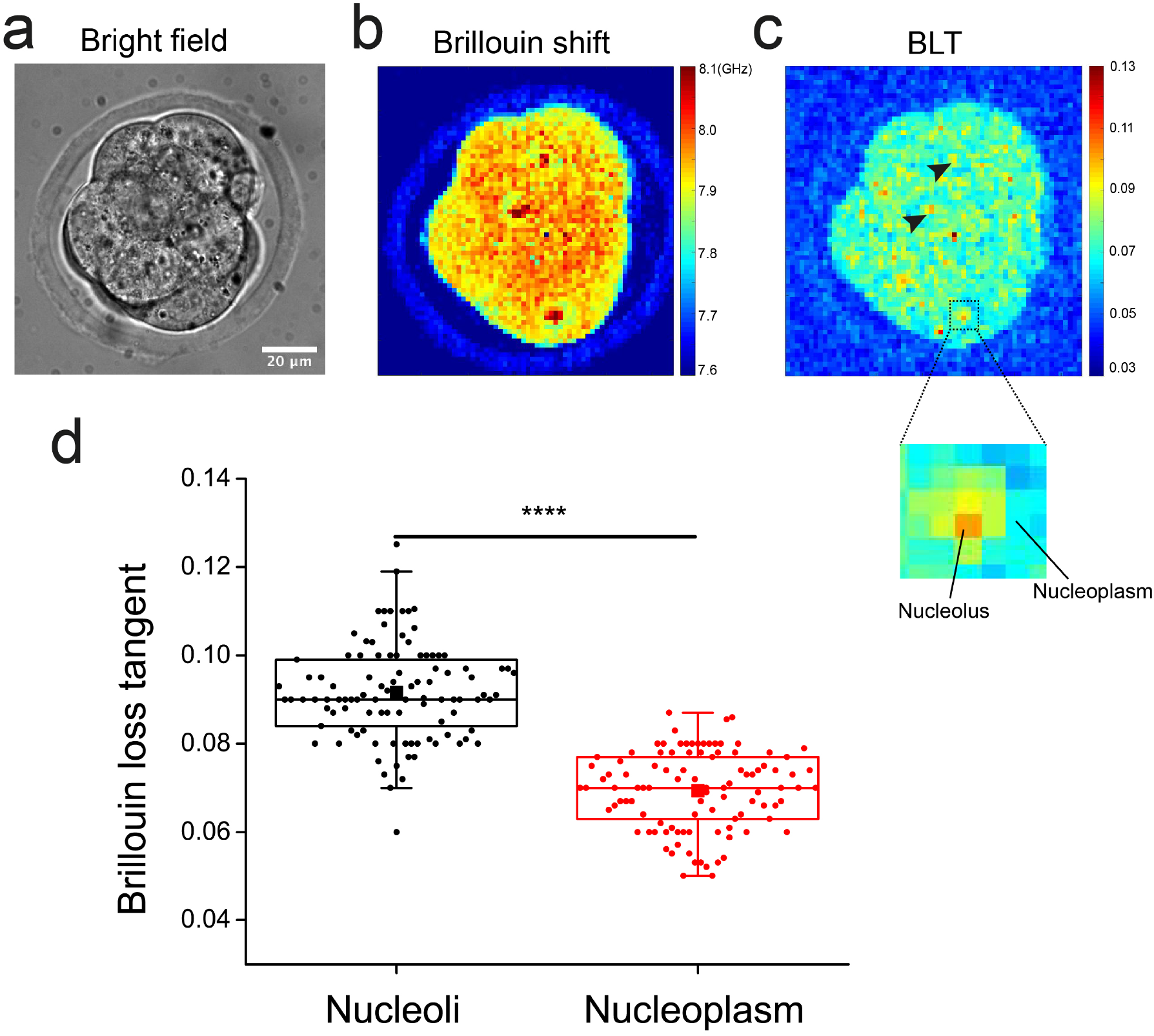
Cell nucleoli have distinctly higher relative viscous components than that of the nucleoplasm. **a.** Brightfield image of an 8-cell stage mouse embryo. **b.** Map of Brillouin shift for the same mouse embryo in **a**, showing clearly the nuclei and nucleoli in three of the blastomeres. c. Corresponding Brillouin loss tangent for (a). Black arrowheads indicate the nucleus region. Box indicates a zoom out sub-nuclear region. **d.** Brillouin loss tangent for nucleoli compared to the nucleoplasm. Data pooled from 102 mouse embryos at various stages of preimplantation. \*\*\*\**P*< 0.0001.

**Supplementary Figure 2:**
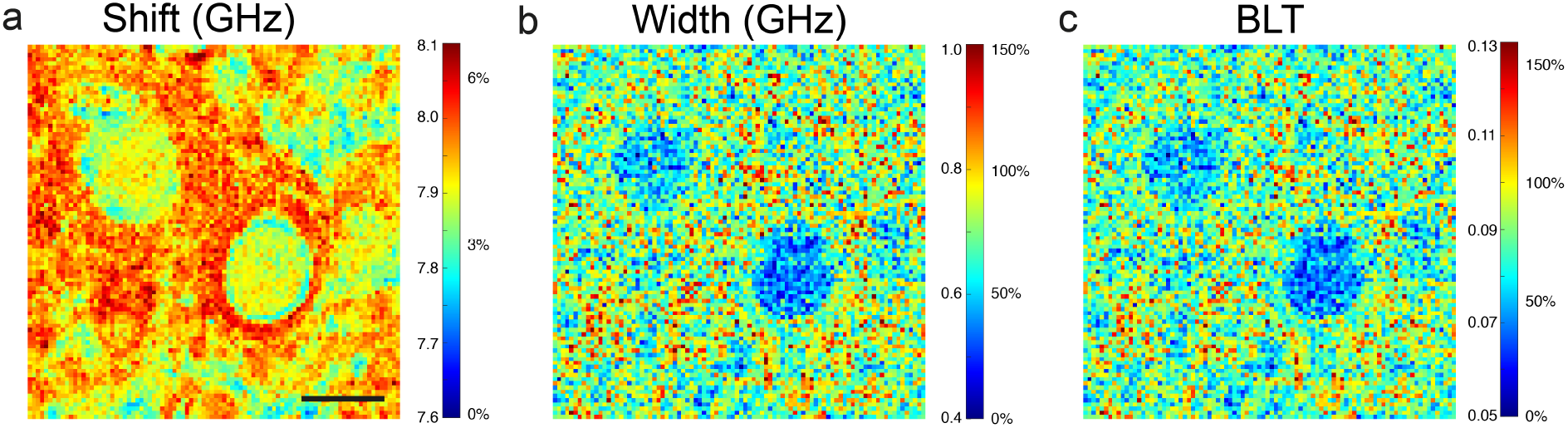
Changes in BLT are mainly due to a change in the Brillouin linewidth and tissue micro-viscosity. Brillouin shift (**a**), width (**b**) and BLT (**c**) for a P7 ovarian interior (**Fig. 1a** in the main texts). Scale bar = 40 μm. Heat map bars indicate absolute values of the shift and width (left side of the bar) and percentage change relative to the medium (7.6 GHz and 0.4 GHz, right side of the bar). The relative change in BLT is approximately the sum of the relative changes of the shift and width. We therefore infer that the main contribution to changes in BLT is the linewidth and hence the effective micro-viscosity of the tissue.

**Supplementary Figure 3:**
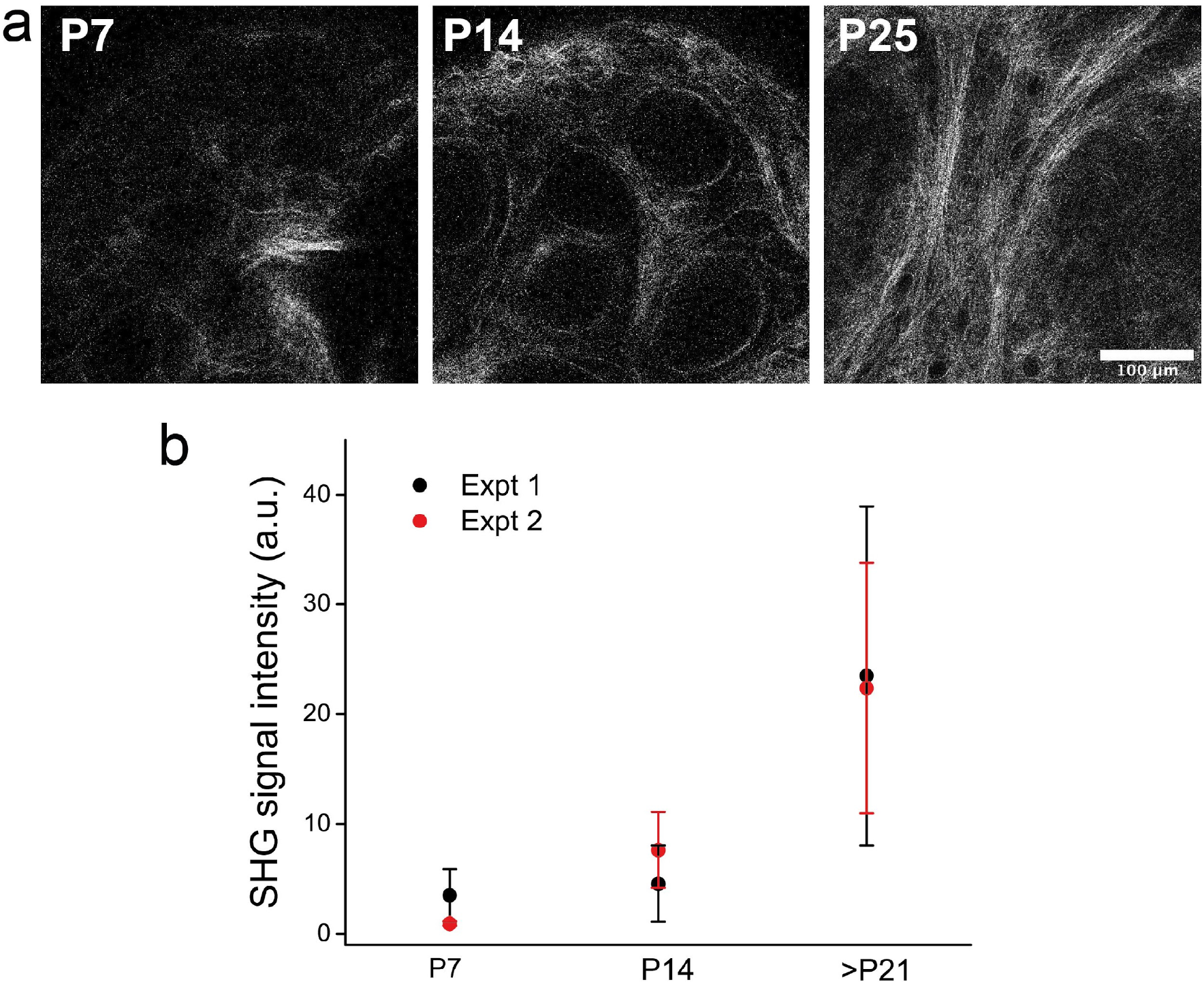
Second Harmonic Generation imaging of fibrillar collagen deposition in mouse ovaries of various stages of folliculogenesis. **a.** Representative images of P7, P14 and P25 mouse ovaries. **b.** Plot of interstitial SGH signal intensity with follicle maturation, indicating that the interstitial collagen network becomes denser and more fibrillar during follicle maturation. PF: Primordial follicle, SF: Secondary follicle, AF: Antral follicle. *N* = 9 from four P7 ovaries, *N* = 17 from four P14 ovaries, and *N* = 23 from four P25 ovaries.

